# A systems-level machine learning approach uncovers therapeutic targets in clear cell renal cell carcinoma

**DOI:** 10.1101/2025.05.26.656158

**Authors:** Silas Ruhrberg Estévez, Greta Baltusyte, Gehad Youssef, Namshik Han

## Abstract

We present a generalisable, interpretable machine learning framework for therapeutic target discovery using single-cell transcriptomics, protein interaction networks, and drug proximity analysis. The pipeline integrates feature selection via gradient boosting classifiers, systems-level network inference, and in silico drug repurposing, enabling the identification of actionable targets with cellular specificity. As a proof of concept, we apply the method to clear cell renal cell carcinoma (ccRCC), an aggressive kidney cancer with limited treatment options. The model identifies 96 tumour-intrinsic genes, refines them to 16 targets through CRISPR screens and biological curation, and prioritises FDA-approved compounds via network-based proximity scoring. Several novel therapeutic mechanisms - including ABL1, CDK4/6, and JAK inhibition - emerge from this analysis, with predicted compounds showing superior efficacy to standard-of-care drugs across multiple ccRCC cell lines. Beyond ccRCC, this framework offers a scalable strategy for drug discovery across diverse diseases, combining machine learning interpretability with systems biology to accelerate therapeutic development.

## Introduction

Each year, kidney cancer is diagnosed in over 400,000 individuals worldwide, with global incidence having doubled over the past three decades^1 2^. The most common subtype, clear cell renal cell carcinoma (ccRCC), arises from the renal epithelium and accounts for approximately 75% of cases^3^. ccRCC is characterised by high metastatic potential and poor prognosis׈. While partial or radical nephrectomy is effective for localised disease׈, around 20% of patients present with metastases at diagnosis, and a further 30% relapse after surgery׈. Treatment of metastatic ccRCC remains challenging due to limited responsiveness to systemic therapies and a five-year survival rate below 10%׈. These poor outcomes, coupled with rapid development of resistance to current treatments, highlight the need for data-driven, computational frameworks that can identify novel therapeutic targets beyond current clinical strategies.

Currently, 16 FDA-approved drugs are available for the treatment of renal cell carcinoma, grouped into four major categories׈. Immunotherapies such as Ipilimumab, Avelumab, Pembrolizumab, Nivolumab, and Aldesleukin enhance tumour recognition by modulating immune checkpoints (e.g., CTLA-4, PD-L1) or stimulating T-cell proliferation via IL-2. Angiogenesis inhibitors - including Bevacizumab, Axitinib, Belzutifan, Tivozanib, Sorafenib, and Sunitinib - suppress tumour vascularisation by disrupting VEGF and HIF-2α signalling pathways. The mTOR inhibitors Everolimus and Temsirolimus target tumour growth via mTOR pathway inhibition. More recently, multitarget kinase inhibitors such as Cabozantinib and Lenvatinib have emerged to simultaneously inhibit VEGF receptors alongside additional tumour-promoting kinases^3^.

Despite the availability of multiple therapeutic agents, treatment responses in ccRCC remain suboptimal. Resistance to targeted therapies typically develops within 8–9 months of first-line treatment and 5–6 months for second-line regimens^3^. Non-targeted agents also yield limited benefit: the response rate for Nivolumab is approximately 25%, while Everolimus achieves only 5%׈. Traditional chemotherapies such as Vinblastine and 5-fluorouracil are rarely used due to poor efficacy (<10%) and nephrotoxicity^1^׈^11^. Notably, most approved therapies act through modulation of the immune or vascular microenvironment, rather than addressing tumour-intrinsic molecular vulnerabilities. This highlights the pressing need for systematic, data-driven approaches that can uncover cell-intrinsic therapeutic targets with greater precision.

Several efforts have sought to address these therapeutic limitations by repurposing compounds that target molecular markers identified through genomic data^12 13^. Drug repurposing offers an efficient development path, leveraging compounds with established safety profiles and mechanisms of action^1^׈. Prior approaches for ccRCC have primarily relied on bulk RNA sequencing to identify differentially expressed genes, or on isolated target discovery using single-cell RNA sequencing (scRNA-seq). However, these methods often lack external validation and fail to model the regulatory complexity and transcriptional heterogeneity captured at the single-cell level. Moreover, existing machine learning approaches to drug repurposing often rely on black-box models that limit interpretability, making it difficult to trace predictions back to biological mechanisms. Furthermore, they typically disregard protein–protein interaction networks and systems-level processes critical to tumour progression. This underscores the need for integrative, interpretable computational frameworks that can leverage single-cell resolution data to uncover robust, tumour-intrinsic drug targets.

We present a systems-level machine learning framework for drug target discovery that integrates single-cell transcriptomics, interpretable feature selection, protein interaction networks, and network-based drug proximity analysis. Applied to clear cell renal cell carcinoma (ccRCC) as a proof of concept, the pipeline identifies five therapeutic mechanisms not currently employed in standard care, several of which demonstrate efficacy in preclinical validation. In contrast to prior approaches that rely on bulk transcriptomic data or single-gene markers, our method leverages public scRNA-seq datasets for high-resolution modelling of tumour-intrinsic vulnerabilities and enables rapid in silico validation across independent cohorts. By combining cellular-resolution expression data with network-based inference, the framework captures dynamic regulatory mechanisms underlying disease progression. It not only uncovers novel therapeutic opportunities but also recovers clinically established targets, underscoring its robustness. While validated here in the context of ccRCC, the pipeline is modular and generalisable, offering a scalable strategy for AI-guided drug repurposing discovery across diverse cancer types and disease contexts.

By integrating explainable machine learning with multiscale biological data, our approach contributes a practical framework for interpretable AI in biomedical discovery - bridging computational prediction and translational relevance.

## Methods

### Single cell RNA data preprocessing

The training dataset was obtained from a single centre and comprised samples from ten ccRCC tumours, one oncocytoma, and one benign thick-walled cyst^1^׈. Cells were collected from normal kidney tissue, multiple tumour regions, and peripheral blood. Each cell was individually sequenced and subjected to quality control. Clustering and cell type annotation were performed using the *Seurat* package in R. The final dataset included over 270,000 cells.

For validation, we utilized six publicly available datasets:

1. Three single-cell RNA sequencing (scRNA-seq) datasets from tumour and adjacent tissues, originating from studies conducted in the United States^1^׈, China^1^׈, and Lithuania^1^׈.
2. One scRNA-seq dataset from the United States focusing on bone metastases and adjacent healthy tissue^1^׈.
3. Two bulk RNA-seq datasets^2^ ^21^׈.

For each dataset, we used either the published cell annotations or re-annotated cells using the *Seurat* pipeline, applying marker genes provided in the respective studies. Although cell type labelling varied across datasets, this did not impact our analysis, as we focused primarily on distinguishing tumour cells from non-tumour cells. Donor annotations were available for all bulk RNA-seq samples. Genes absent in single-cell validation datasets were imputed as zero, while genes missing in the bulk RNA-seq datasets were excluded from the analysis, as they did not influence UMAP representations.

### Drug target identification

All subsequent data analysis was conducted using Python 3.12.8. For initial model exploration and classifier selection, we applied an 80:20 train–validation split stratified by cell type, for both multiclass and binary classification tasks. The final model was trained on the full dataset and evaluated on external test sets. Feature selection was performed using five-fold cross-validation, also stratified by cell type. As the binary tumour classification task was highly imbalanced, we applied the Synthetic Minority Oversampling Technique (SMOTE) to increase the representation of tumour cells in the training data^22^.

Both classification and feature selection were performed using the LightGBM algorithm from the Python package *lightgbm*^23^. Univariate feature selection was conducted using the *feature-engine* library, in combination with a LightGBM model consisting of 1,000 estimators, a maximum tree depth of 10, and 31 leaves, trained with a learning rate of 0.05 to minimize binary log loss. Feature performance was assessed by computing the average area under the receiver operating characteristic curve (AUC). For final classification, a more robust LightGBM model was used with 10,000 estimators and a learning rate of 0.01.

All UMAP visualizations were generated using the Python package *umap*, with the number of neighbours set to 15. The classifier’s performance on external datasets was assessed using accuracy, precision, recall, F1 score, and ROC-AUC to evaluate robustness across multiple performance metrics^2^׈.

The identified drug targets were mapped to their corresponding proteins using BioMart. A protein–protein interaction (PPI) network was constructed using the *igraph* library, with interaction data obtained from the STRING database^2^׈, applying a confidence threshold of 0.425.

We previously developed a network-based drug repurposing pipeline to identify FDA-approved compounds targeting SARS-CoV-2 pathways^2^׈. In the present study, we applied the same network proximity analysis method, as described in Han et al., to assess the therapeutic relevance of key proteins identified within the ccRCC-specific interaction network. Specifically, given a set of key proteins KKK from our PPI network and a set of known drug targets TTT, we computed network proximity using the “closest” distance metric outlined by Guney et al.^2^׈, which has demonstrated effectiveness in drug–disease prediction tasks. The shortest distances between proteins were calculated using the *igraph* implementation of Dijkstra’s algorithm.

To evaluate the significance of the observed proximities, we performed 1,000 permutations and calculated associated z-scores. A binning strategy was used during random sampling to preserve the node degree distribution of the original network. Proteins with a proximity z-score below –2 (p < 0.01) were considered significantly proximal to known drug targets and prioritized as candidate therapeutic targets^2^׈.

To refine the target list, proteins classified as “common essentials” in the DEPMAP CRISPR 24Q4 screen were excluded^2^׈. The remaining candidates underwent manual curation to assess biological plausibility. Protein localization, function, and prognostic relevance in ccRCC were evaluated using The Human Protein Atlas^2^׈.

### Hit identification

We screened all FDA-approved drugs from DrugBank and non-FDA-approved compounds from PubChem. For each compound, network proximity to key proteins was calculated, and candidates with z-proximity scores below –2 were prioritized. The resulting drug list was then reviewed to ensure favourable safety profiles and known anti-cancer activity, while excluding compounds previously applied to renal cell carcinoma. To verify novelty, we searched ClinicalTrials.gov and PubMed using combinations of the drug name with the terms “renal carcinoma” or “kidney cancer.” Finally, drugs were categorized by their mechanisms of action to identify the most suitable compound for each therapeutic strategy.

### Preclinical drug validation

We assessed the efficacy of selected compounds using the PRISM Repurposing Public 24Q2 dataset provided by DEPMAP²׈. Our analysis focused on FDA-approved drugs predicted to reduce tumour cell viability based on their mechanisms of action. To maintain biological relevance, agents with broad systemic effects, such as immune checkpoint inhibitors and corticosteroids, were excluded.

The DEPMAP screen included 20 renal cell carcinoma (RCC) cell lines, of which three were excluded due to their classification as non-ccRCC subtypes. Compounds were tested using the PRISM assay at a concentration of 2.5μM for five days. Each compound was screened in triplicate, with every plate containing positive controls (bortezomib, 20׈μM) and negative controls (DMSO) to ensure assay consistency. Differences in cell viability were evaluated using a two-tailed Mann–Whitney U test. A cell line was considered responsive if the log fold change in viability was below zero.

## Results

### Drug target selection using machine learning

We developed a novel drug discovery pipeline that integrates single-cell RNA sequencing (scRNA-seq), machine learning, and network analysis (see Figure׈1A). The training dataset comprised over 270,000 cells collected from ten ccRCC tumours across multiple anatomical sites, as well as from adjacent healthy tissue samples^1^׈. In total, the dataset spans nine distinct cell types, as visualized in Figure 1B & C.

**Figure 1.**
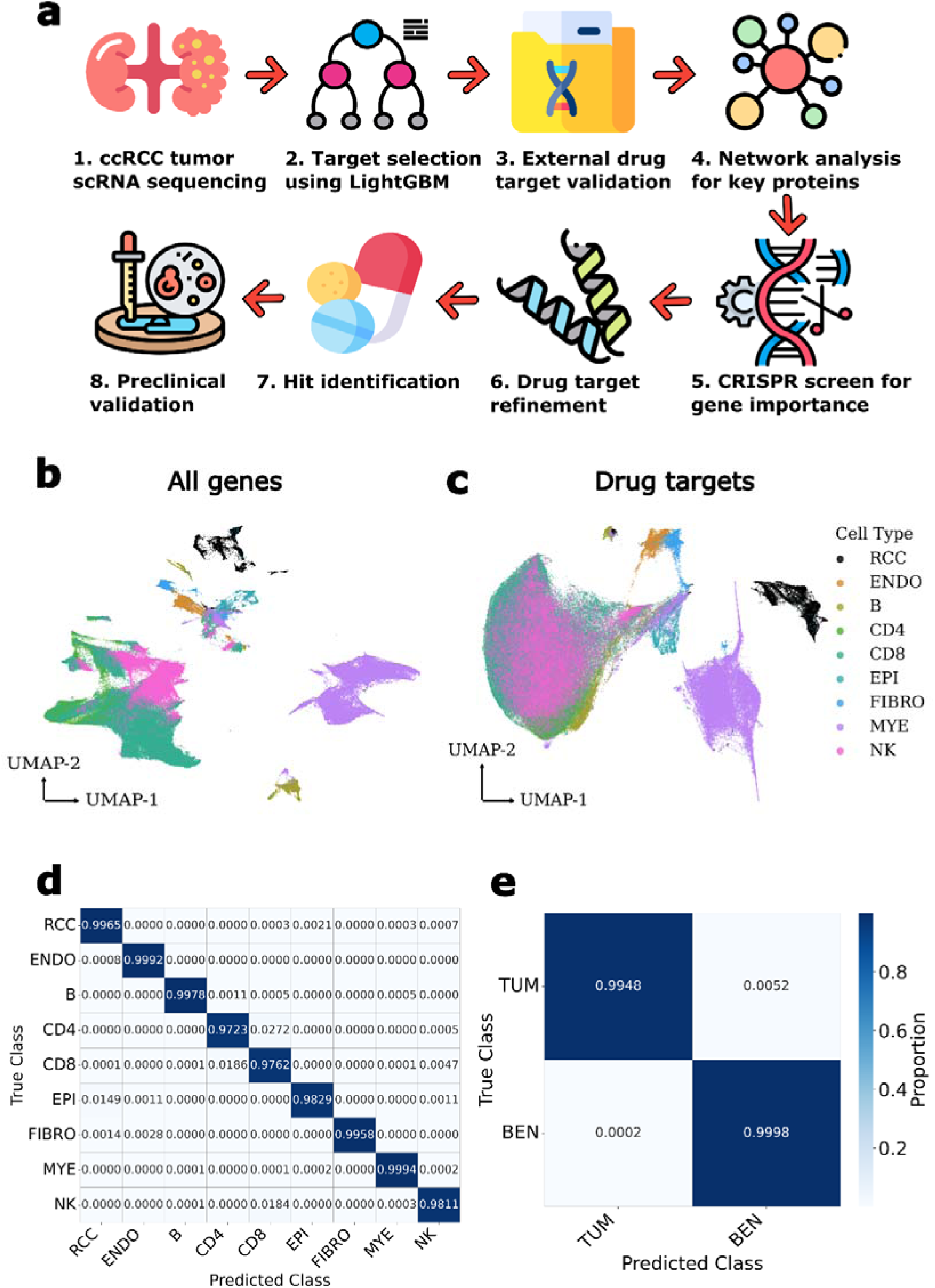
Integrative machine learning pipeline identifies tumour-specific gene signatures from single-cell RNA-seq data. **a**) Overview of the systems-level drug discovery pipeline integrating scRNA-seq, LightGBM-based classification, protein network analysis, and drug proximity scoring to identify repurposable therapeutics for ccRCC. **b**–**c**) UMAP visualizations of the training dataset showing cell type clusters (**b**) and tumour–normal separation based on the selected 96-gene feature subset (**c**), defined as genes with AUC > 0.8. **d**) Confusion matrix of the multiclass classifier distinguishing nine major kidney cell types on validation data. **e**) Confusion matrix of the binary classifier distinguishing tumour from non-tumour cells in validation data.

To validate cell-type specificity, we trained both a multiclass LightGBM classifier to distinguish all cell types and a binary classifier to separate tumour from non-tumour cells^23^. Both models demonstrated high accuracy on held-out data (see Figure׈1D & E), indicating that the LightGBM architecture is well suited for capturing transcriptional differences in scRNA-seq data.

Next, we performed univariate feature selection with five-fold cross-validation to identify genes that best discriminated tumour cells from normal cells. Using a threshold of area under the receiver operating characteristic curve (AUC) ≥L0.8, we identified 96 genes as potential drug targets (see Supplementary Table׈1). The distribution of AUC values for these genes is presented in Supplementary Figure׈1A. A full list of gene names and their corresponding AUC scores is available in Supplementary File׈1, while log fold changes in expression between tumour and non-tumour cells are provided in Supplementary File׈2.

### Drug target characterization

Analysis of drug target localization and protein translation patterns revealed that the selected targets tended to have fewer exons than average (see Supplementary Figure 1E), although they were uniformly distributed across chromosomes (see Supplementary Figure׈1B). Genes with fewer exons are often involved in cell cycle regulation and other rapid-response processes, where reduced splicing complexity facilitates more efficient transcription and translation^3^.

We next examined the biological functions of the drug targets. Using STRING, we constructed a protein–protein interaction (PPI) network, which revealed strong connectivity among the majority of targets (see Figure׈2A) ^31^. Separate analyses of PPI networks for upregulated and downregulated genes further supported the functional interdependence of the identified targets (see Supplementary Figure 1׈C & D).

**Figure 2.**
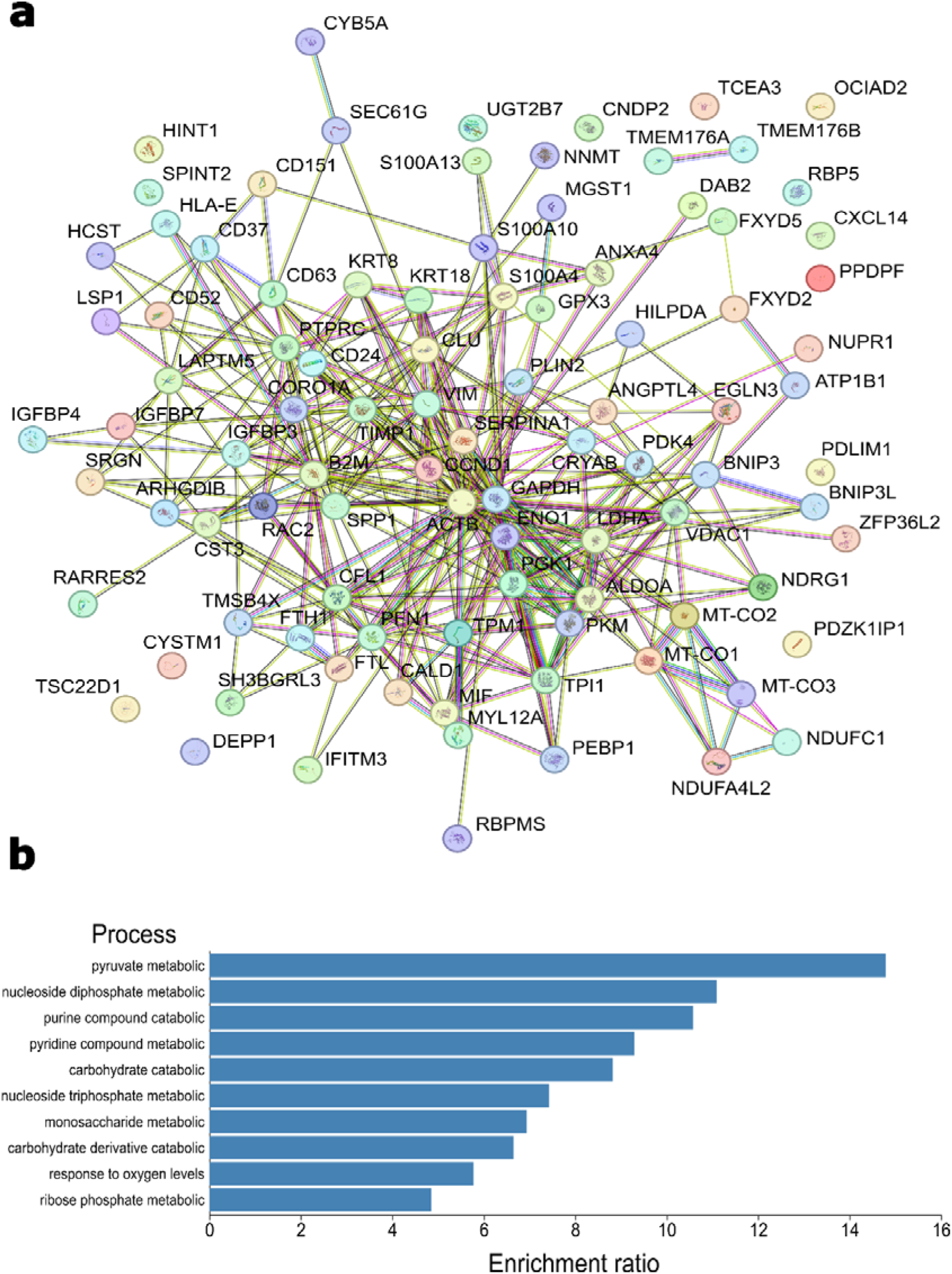
Functional and network characterization of tumour-specific drug targets. **a**) Protein–protein interaction (PPI) network of the 96 selected drug targets, generated using STRING, demonstrating high connectivity and pathway convergence^31^. **b)** Pathway enrichment analysis using WebGestalt reveals that drug targets are significantly enriched in pathways related to metabolism, oxygen sensing, and proliferation. Bar plot shows top enriched pathways ranked by enrichment ratio.

We then explored the biological pathways associated with these proteins. The most significantly enriched pathways included carbohydrate metabolism, nucleoside turnover, and oxygen level responses (see Figure 2B). The enrichment of carbohydrate metabolic pathways is consistent with the well-documented reliance of kidney tumours on glycolysis, characteristic of the Warburg effect^32 33^. Aberrant oxygen regulation, often mediated by VHL mutations, has also been extensively studied in ccRCC^3^׈. Similarly, altered nucleoside metabolism is frequently observed in cancer, where it supports the increased demand for nucleotides during rapid proliferation^3^׈.

To further validate the biological relevance of these targets, we examined their associations with key cancer hallmarks, including apoptosis, angiogenesis, and metabolic reprogramming. Most hallmarks were represented within the target gene set, with many individual targets contributing to multiple hallmark processes (see Supplementary Figure׈2A & B).

### Drug target signature validation

To validate the drug target signature, we utilized six external datasets originating from three different continents. The first dataset, collected in the United States, consists of scRNA-seq data from seven ccRCC tumour specimens and six benign human kidney samples^1^׈. The second, from China, includes scRNA-seq data from seven ccRCC tumours paired with five adjacent normal kidney samples¹L. The third, collected in Lithuania, contains scRNA-seq profiles from eight ccRCC tumours and nine matched healthy-adjacent kidney tissues^1^׈. In addition, we incorporated a fourth scRNA-seq dataset from the United States, comprising nine ccRCC bone metastasis samples^1^׈.

To provide broader population-level validation, we also analysed two bulk RNA-seq datasets. The TCGA ccRCC cohort includes 533 tumour samples and 73 normal kidney samples^2^׈. The second dataset, from Laskar et al., includes 503 tumour samples and 153 normal kidney samples^21^.

We trained a LightGBM (LGBM) model to perform binary classification of tumour versus normal cells using the 96 drug targets across all cells in our training dataset These genes demonstrated strong discriminatory power, as visualized by UMAP embeddings that showed clear separation between tumour and non-tumour populations (Figure 3A–D). The classifier was then applied to predict cell identities in the four external scRNA-seq datasets. The drug targets effectively distinguished cancerous from normal cells in all cases, with the classifier achieving high performance across datasets (see Figure׈3G). The AUC values were as follows: United States (0.99), China (0.99), Lithuania (0.99), and Metastasis (0.92). Additional performance metrics are provided in Supplementary Table׈1.

**Figure 3.**
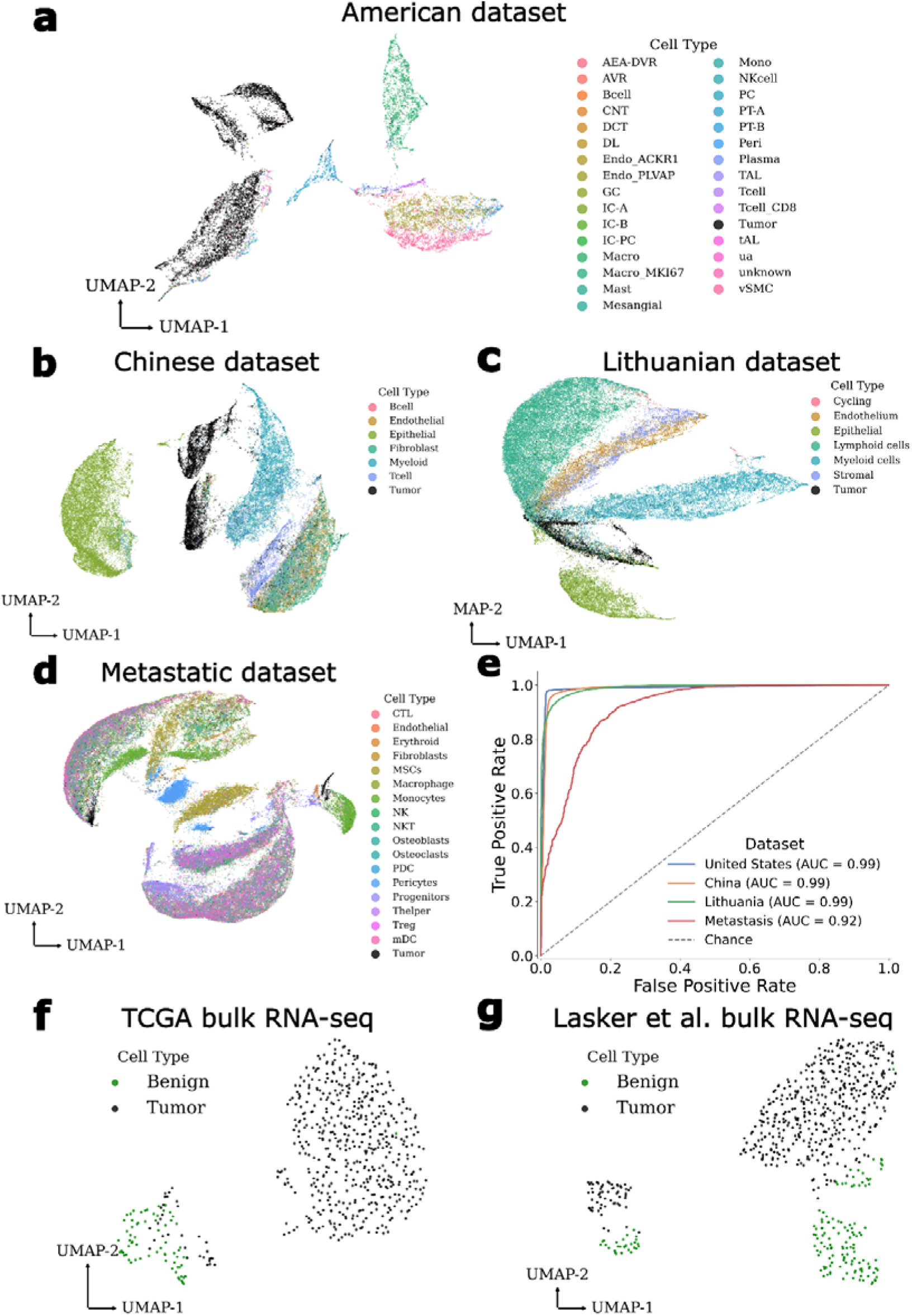
**External validation confirms robustness of tumour-specific gene signature across independent datasets a–d**) UMAP visualizations show effective tumour versus non-tumour separation using the 96-gene feature set (AUC ≥ 0.8) in four scRNA-seq datasets from the United States (Zhang et al. 2021) (**a**), China (Zhang et al. 2022) (**b**), Lithuania (Zvirblyte et al. 2024) (**c**), and ccRCC bone metastases (Mei et al. 2024) (**d**). **e**) ROC curves and AUC values confirm consistent classifier performance across diverse scRNA and bulk cohorts, with AUC values >0.9 in all datasets. **f–g**) UMAP plots of two bulk RNA-seq datasets (TCGA and Laskar et al. 2021) also demonstrate separation of tumour and normal samples despite cellular averaging.

As expected, this model is not compatible with bulk RNA-seq data due to fundamental differences in data modality. Unlike scRNA-seq, bulk RNA-seq averages gene expression across diverse cell populations, potentially masking cell-type-specific signals. For example, a drug target upregulated in infiltrating lymphocytes may appear downregulated in tumour cells at the single-cell level but elevated in bulk tissue due to immune cell presence. Nevertheless, visualization using UMAP showed that the drug targets were able to clearly separate tumour and normal samples even in bulk datasets, demonstrating the robustness of the signature (see Figure׈3E & F).

### Hit identification

Network analysis of the identified drug targets revealed 92 key nodes (see Supplementary File׈3). Among these proteins, 57.6% were intracellular, 35% secreted, and 17.4% membrane-bound (see Figure׈4D-G). Functionally, the proteins were categorized as follows: 39.1% enzymes, 15.2% receptors, 19.6% ligands, 9.8% transcription factors, and 16.3% structural proteins. Expression analysis indicated that 63% were downregulated in tumour cells, while 37% were upregulated. Notably, 46.7% of the targets showed prognostic significance in ccRCC, and in many cases, ccRCC was the only cancer type in which these genes held prognostic value. Several central nodes, such as MYC, TP53, VEGFA, MAPK1, and CASP3, are well-established oncogenic drivers, further reinforcing the robustness of our network-based target selection (see Figure׈4A & B). The complete protein–protein interaction network, including non-differentially expressed nodes, is provided in Supplementary Figure 3A.

**Figure 4.**
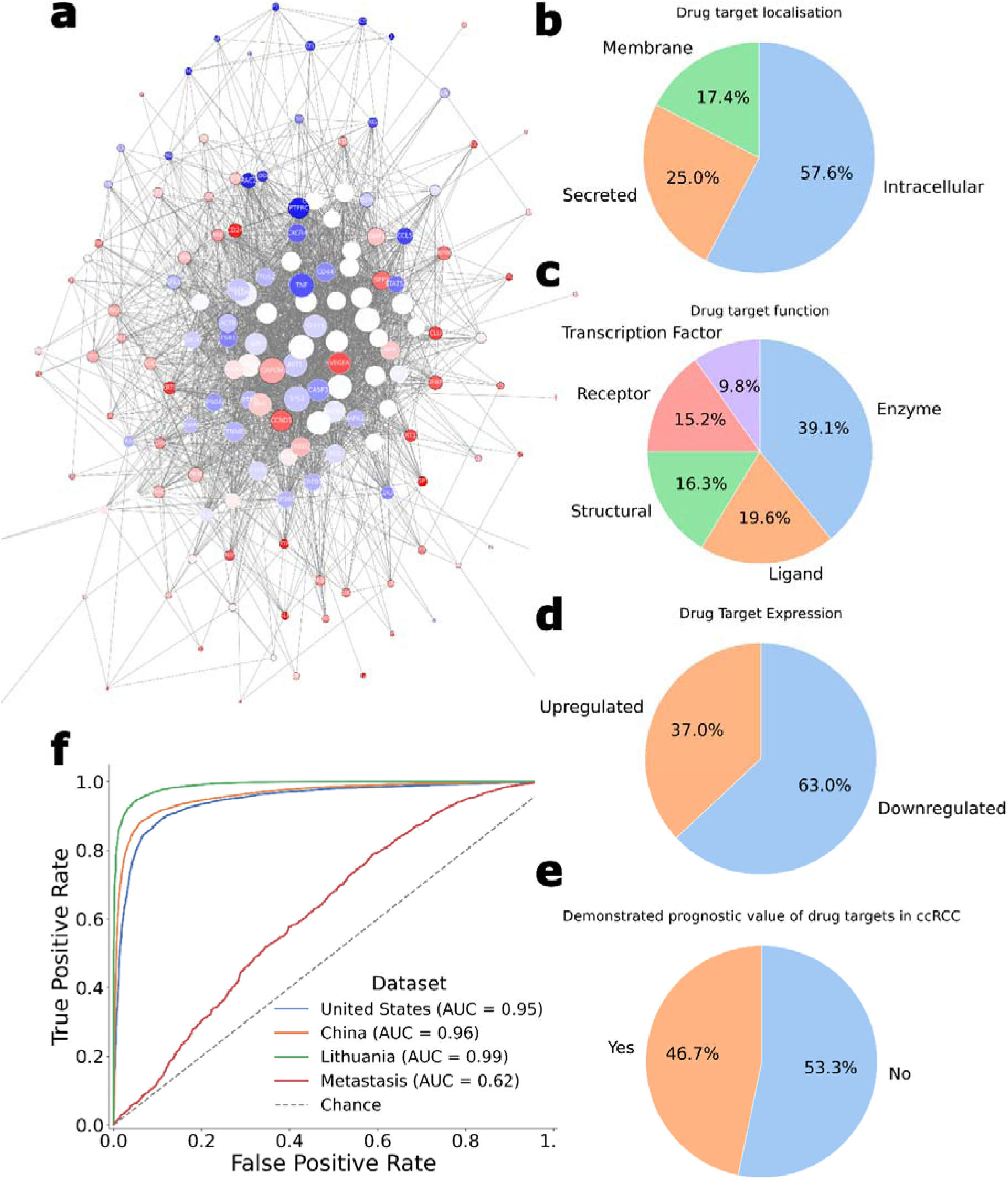
Network-guided refinement of drug targets reveals actionable therapeutic mechanisms. **a**) Drug– target interaction network of the 92 key nodes, organized by subcellular localization: membrane, cytosol, and nucleus. Red = upregulated genes; blue = downregulated genes; white = unchanged. Size of the circles represent the size of the eigenvectors for each key node. **b**) Network highlighting central nodes based on connectivity and functional relevance. **c**) ROC curves showing classifier performance using the refined 16-gene target set across four external datasets. **d–g**) Summary of characteristics of identified key nodes: **d**) Subcellular localization; **e**) Protein function (e.g., enzyme, receptor); **f**) Direction of differential expression in tumour cells; **g**) Prognostic significance in ccRCC based on Human Protein Atlas annotationsAtlas^29^.

To refine this list, we selected drug targets that were both differentially expressed in tumour cells and identified through LightGBM-based feature selection, yielding the intersection of the 92 key nodes and 96 high-AUC genes. To ensure specificity, we excluded genes classified as common essentials by the DEPMAP CRISPR screen^2^. This refinement produced a final list of 23 drug targets (see Supplementary Table׈2). To further refine the drug target set, we manually reviewed published literature for each gene. This step was critical, as some features identified through machine learning may reflect changes in the tumour microenvironment (e.g., immune infiltration) rather than intrinsic tumour biology. Supplementary Table׈2 summarizes the literature review. Seven genes were excluded due to unclear or contradictory links to tumour progression, often showing regulatory patterns inconsistent with prior studies. These were deemed more likely to reflect microenvironmental changes than true oncogenic drivers.

Using the remaining 16 targets, classification performance (AUC) remained high across the four scRNA-seq datasets: United States (0.94), China (0.96), Lithuania (0.91), and Metastasis (0.62) (see Figure׈4C). Additional metrics are presented in Supplementary Table 3. UMAP visualization confirmed that the refined target set still effectively separates distinct cell populations (see Supplementary Figure׈4A-F).

We then performed drug proximity analysis on the 16 refined targets using both FDA-approved and non-FDA-approved compounds. Full results are presented in Supplementary Files׈4 and 5. A proximity z-score cutoff of –2 was used to prioritize drug–target associations (see Supplementary Figure׈3B & C). Notably, three approved ccRCC therapies, sorafenib, sunitinib, and pazopanib, were recovered by our pipeline, with sorafenib and sunitinib ranking among the top ten hits. Both pazopanib and sunitinib are first-line treatments for advanced ccRCC^32^, further validating our approach. We also identified sirolimus, a close analogue of everolimus, as a candidate mTOR inhibitor. Several additional VEGFR- and kinase-targeting drugs, such as vandetanib, were also detected.

To prioritize translational potential, we focused on FDA-approved compounds, given their well-characterized safety profiles. This yielded 39 candidate drugs, which we categorized by mechanism of action (see Table׈1). Several of these pathways have known anti-ccRCC activity. Among the previously underexplored mechanisms, we identified ABL1 inhibition, CDK4/6 inhibition, JAK inhibition, and non-VEGFR kinase inhibition as promising therapeutic strategies. Additionally, we identified the non-FDA-approved ROCK inhibitor Fasudil, currently used for subarachnoid haemorrhage, as a potential candidate for ccRCC. However, our downstream analysis focused on FDA-approved compounds due to their greater clinical tractability.

**Table 1.** Therapeutic mechanisms of FDA-approved drugs identified through network-based proximity analysis. Summary of repurposable FDA-approved drugs prioritized using network proximity to ccRCC-specific targets. Compounds are grouped by mechanism of action, with context provided on their relevance to clear cell renal cell carcinoma. Several pathways, such as VEGFR inhibition, mTOR inhibition, and multikinase targeting, are already used clinically, validating the approach. Others, including ABL1, CDK4/6, and JAK inhibition, represent novel strategies with demonstrated preclinical efficacy in this study. Agents with limited renal tolerability or unclear oncological relevance were deprioritized for downstream validation. Drugs indicated in bold very included in the preclinical validation.

| **Mechanism of action (MOA)** | **Drugs identified** | **MOA Comments** |
| --- | --- | --- |
| **VEGFR inhibitors** | Sunitinib, Sorafenib, Pazopanib, Vandetanib, Regorafenib | Already used in clinic |
| **mTOR inhibitors** | Sirolimus | Already used in clinic |
| **JAK inhibitors** | **Tofacitinib, Ruxolitinib** | Promising target |
| EGFR inhibitor | Erlotinib, Gefitinib, Trastuzumab, Afatinib | Successfully trailed in RCC<sup>44</sup> |
| MEK inhibitors | Trametinib | Successfully trailed in RCC<sup>45</sup> |
| **CDK 4/6 inhibitors** | **Ribociclib** | Promising target |
| Calcium metabolism modifiers | Amiodarone, Calcitriol | Benefits unclear |
| **Hormone analogues** | Stanozol, Nilutamide, Fulvestrant, Trilostane, Liotrix | Suggested as treatment for RCC but side-effects<sup>46</sup> |
| Dietary supplements | Zinc chloride | Benefits unclear |
| **ABL1 inhibitors** | **Bosutinib, Dasatinib, Nilotinib, Imatinib, Ponatinib** | Promising target |
| Antibiotic | Gentamicin | Nephrotoxic |
| Cytotoxic | Paclitaxel, Ingenol mebutate | Nephrotoxic |
| Clotting cascade modifier | Tranexemic acid, Tenecteplase | Benefits unclear |
| Glucocorticoids | Hydrocortisone acetate, Hydrocortisone | Successfully trailed in RCC in vitro<sup>47</sup> |
| **Kinase inhibitors** | **Fostamatinib, Encorafenib, Zanubrutinib, Crizotinib** | Promising target |

### Preclinical drug validation

We assessed the efficacy of candidate compounds using the DEPMAP PRISM screen on 17 renal cell carcinoma (RCC) cell lines. As a baseline, we assessed nine FDA-approved therapies currently used to treat clear cell renal cell carcinoma (ccRCC). Consistent with clinical variability in treatment response, none of these agents reduced cell viability in more than 70% of the tested cell lines (see Figure׈5A & C). The most effective among them, the multikinase inhibitor Cabozantinib, achieved a mean logL viability reduction of –0.22, but failed to inhibit approximately 40% of RCC cell lines, consistent with clinical observations of resistance to current treatments.

**Figure 5.**
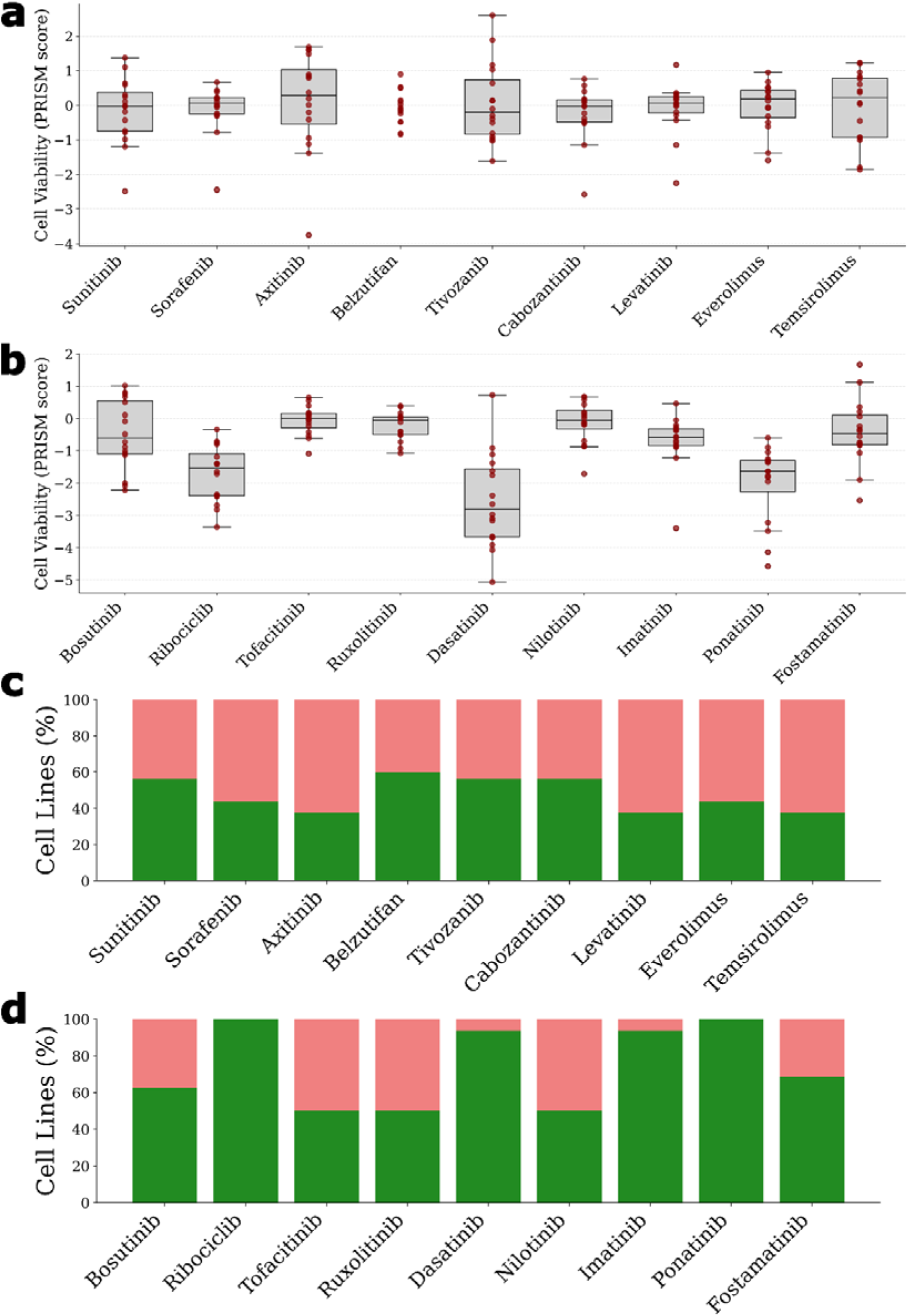
Repurposed drug candidates demonstrate superior efficacy in reducing cancer cell viability compared to standard-of-care therapies. (a–b) LogL fold change in viability of ccRCC cancer cell lines treated with **(a)** approved clinical therapies and **(b)** top-ranked repurposed candidates prioritised by our computational framework, using PRISM 24Q2 screening data. **(c–d)** Proportion of ccRCC cancer cell lines responsive to each compound, defined by drug-induced reduction in cancer cell viability. Repurposed agents - including Ribociclib, Ponatinib, and Dasatinib - exhibited broader efficacy and greater inhibition than all currently approved treatments. Green bars indicate responsive cancer cell lines; pink bars denote non-responders.

In contrast, all nine of our proposed drug candidates demonstrated inhibitory effects comparable to or greater than those of standard-of-care therapies (see Figure׈5B & D). Notably, three predicted candidates, Ribociclib, Ponatinib, and Dasatinib, significantly reduced cell viability, with p-values of 0.00006, 0.00002, and 0.00008, respectively, compared to cabozantinib. The corresponding logL fold changes in viability were –1.7, –2.0, and –2.5, respectively. Furthermore, both Ribociclib and Ponatinib inhibited viability in all tested cell lines, underscoring their potential as effective therapeutic options for RCC.

## Discussion

We introduce a novel computational framework for drug discovery that integrates single-cell RNA sequencing (scRNA-seq), interpretable machine learning, and network-based inference. Applied to clear cell renal cell carcinoma (ccRCC) as a proof of concept, the pipeline addresses key limitations of previous computational drug repurposing approaches. Prior studies in kidney cancer have primarily focused on identifying individual targets^12^ or candidate compounds from differential expression in bulk RNA sequencing data^3^׈. Although some were validated experimentally, bulk transcriptomic methods obscure cell-type-specific signals critical to tumour-specific targeting. Other systems-level efforts have also relied on bulk RNA-seq to infer potential targets^13^, limiting their resolution and adaptability. In contrast, our pipeline leverages scRNA-seq data for cell-type-resolved target discovery and incorporates robust validation across independent datasets, offering a scalable, generalisable approach for interpretable AI in precision oncology.

In contrast to prior approaches, our pipeline integrates cell-type-specific transcriptional data with network-based drug proximity analysis to enable precise, interpretable identification of tumour-intrinsic vulnerabilities. While experimental drug screening - conducted without sequencing data - remains a complementary route for drug repurposing^3^׈, such methods are labour-intensive, susceptible to off-target effects, and often lack scalability across disease models. By leveraging single-cell data and systems-level modelling, our computational framework offers a more efficient and generalisable alternative for prioritising clinically actionable targets.

Importantly, prior computational studies have rarely validated their predictions on external datasets, despite the growing availability of such public data. Most also failed to recover drugs currently used in clinical practice - an omission that limits clinical relevance. While this is understandable in studies focused on isolated gene mechanisms, it raises concerns for more comprehensive approaches that rely on bulk RNA signatures, which may be confounded by signals from the tumour microenvironment. In contrast, our framework demonstrates robust performance across independent scRNA-seq and bulk datasets and successfully rediscovers known therapeutic mechanisms, reinforcing its biological validity and translational potential.

Our analysis recovered several drug mechanisms aligned with current ccRCC treatments, including VEGF receptor-targeting kinase inhibitors and multitarget kinase inhibitors. mTOR inhibitors also emerged, though at lower ranks, consistent with their modest clinical efficacy in ccRCC. The ability of the pipeline to rediscover established therapeutic pathways underscores both the robustness of the feature selection strategy and the biological relevance of the inferred targets. This validation highlights the model’s potential not only for novel discovery but also for reliably identifying known tumour progression drivers.

Our analysis also surfaced several compounds that are synthetic derivatives of endogenous hormones. These included the androgen Stanozolol and the antiandrogen Nilutamide - agents used in prostate cancer and performance enhancement. Although their therapeutic relevance in ccRCC remains uncertain, their emergence aligns with epidemiological data indicating a nearly two-fold higher incidence of ccRCC in males^3^׈. Similarly, the oestrogen receptor antagonist Fulvestrant and the steroidogenesis inhibitor Trilostane, both used in breast cancer, were also identified. Oestrogen is known to play a protective role in ccRCC development^3^׈. The presence of drugs with opposing hormonal effects reflects the undirected nature of proximity-based network scoring, which ranks compounds by mechanistic proximity rather than assumed therapeutic direction. Hormonal agents such as Medroxyprogesterone have previously been trialled in ccRCC but were largely discontinued due to side effects and limited efficacy.

In the hit identification stage, we prioritized drug mechanisms that could complement existing treatments to address resistance in ccRCC. Our pipeline identified four underexplored strategies using FDA-approved compounds: ABL1 inhibition, CDK4/6 inhibition, Janus kinase inhibition, and non-VEGFR kinase inhibition. To improve clinical tractability, we excluded agents with broad systemic effects, such as corticosteroids, and filtered out nephrotoxic compounds. From this filtered pool, nine candidates representing these mechanisms (see Table׈1) were selected for preclinical evaluation alongside nine standard-of-care therapies (Figure׈5A–B). All selected candidates demonstrated tumour cell viability inhibition comparable to or exceeding current treatments. Notably, Ponatinib and Dasatinib (ABL1 inhibitors) and Ribociclib (a CDK4/6 inhibitor) significantly outperformed Cabozantinib - the most effective clinically approved agent in our assay - highlighting their potential as viable alternatives or combination partners.

To explore combination potential, we evaluated pharmacological interactions using the DrugBank interactions database. Dasatinib was identified as a CYP3A4 inhibitor, which may compromise the metabolism of frontline agents such as Pazopanib, reducing its suitability for combination. In contrast, both Ponatinib and Ribociclib were predicted to be pharmacologically compatible with Pazopanib, enabling the potential for combination therapies targeting three distinct pathways. Ponatinib, approved for leukaemia, primarily inhibits BCR–ABL but also targets KIT, RET, and Src kinases^1^׈. Ribociclib, a CDK4/6 inhibitor used in breast cancer, has previously been proposed for repurposing in solid tumours, including RCCL^2^׈. These findings highlight the translational value of integrating drug interaction knowledge with network-informed target discovery.

We present a novel computational framework for drug discovery that integrates single-cell transcriptomics, machine learning, and network-based inference. The pipeline enables interpretable identification of therapeutic targets, which can be externally validated across independent datasets. In our ccRCC case study, the framework accurately recovered standard-of-care drug mechanisms and prioritised three FDA-approved compounds - Ponatinib, Dasatinib, and Ribociclib - with distinct mechanisms of action and superior preclinical efficacy. Given the high resistance rates associated with existing therapies, these compounds hold promise for improving clinical outcomes, particularly as components of combination strategies. More broadly, the pipeline offers a scalable and generalisable approach for AI-guided therapeutic discovery across diverse disease contexts.

This study has several limitations. First, while the prioritised compounds have demonstrated safety and anti-cancer activity in other tumour types, they have not yet been evaluated clinically in the context of ccRCC. Future work will involve in vivo and clinical studies to assess their efficacy in this setting. Second, the network proximity metric employed is undirected, which may result in the selection of targets that are biologically proximal but not directly druggable, requiring manual curation to refine the candidate list. Third, the pipeline primarily leverages transcriptional differences between tumour and healthy cells, which may underrepresent the role of the tumour microenvironment. Integrating datasets with richer cell-type diversity from normal kidney tissue and incorporating spatial or multi-omics data could enhance the identification of targets related to immune evasion and stromal signalling.

Beyond its application to ccRCC, this study contributes a modular and reusable framework for integrating single-cell transcriptomic data with interpretable machine learning and network-based drug discovery. Each component - from classifier training to feature selection and proximity scoring - is model-agnostic and adaptable, supporting deployment across diverse cancer types and disease settings. As single-cell and multi-omics datasets continue to expand, this framework offers a scalable foundation for AI-driven therapeutic discovery in biologically complex conditions, particularly where treatment resistance or toxicity limits current options. More broadly, it advances the development of transparent and generalisable machine learning approaches in precision medicine.

## Contributions

S.R.E., G.Y., and G.B. conducted the data analysis. G.Y. conceptualized the project. S.R.E. drafted the first version of the manuscript. G.Y. and N.H. provided overall guidance. All authors reviewed the manuscript.

## Data sharing statement

All datasets used in this study are publicly available. The scRNA training data can be downloaded from Mendeley Data at: https://doi.org/10.17632/g67bkbnhhg.1. The scRNA validation data is accessible via the Gene Expression Omnibus (GEO) repository (https://www.ncbi.nlm.nih.gov/geo/) under accession numbers GSE156632, GSE159115, GSE242299, and GSE202813. The TCGA bulk RNA dataset can be downloaded from their website (https://xenabrowser.net/datapages/). The other bulk RNA-seq dataset is available in the GEO repository under accession number GSE242299. The underlying code for this study is available on GitHub and can be accessed via this link https://github.com/Sr933/rcc.git. The code can also be requested from Silas Ruhrberg Estévez.

## Acknowledgements

We would like to thank Professor Grant Stewart and Dr James Jones (Department of Surgery, University of Cambridge) for their guidance. Figures were generated with icons from Flaticon.

## Role of the funding body

The funder played no role in study design, data collection, analysis, interpretation and writing of the manuscript.

## Ethics

Since both the bulk RNA-seq and scRNA-seq datasets, as well as the preclinical cell viability data, were derived from previously published studies, no ethical concerns were raised.

## References

1. Bukovina, L. et al. Epidemiology of Renal Cell Carcinoma: 2022 Update. Eur. Urol. 82, 529–542 (2022).

2. Du, Z., Chen, W., Xia, Q., Shi, O. & Chen, Q. Trends and projections of kidney cancer incidence at the global and national levels, 1990-2030: a Bayesian age-period-cohort modelling study. Biomark. Res. 8, 16 (2020).

3. Hsieh, J. J. et al. Renal cell carcinoma. Nat. Rev. Dis. Primer 3, 17009 (2017).

4. Schiavoni, V. et al. Recent Advances in the Management of Clear Cell Renal Cell Carcinoma: Novel Biomarkers and Targeted Therapies. Cancers 15, 3207 (2023).

5. Mattila, K. E., Vainio, P. & Jaakkola, P. M. Prognostic Factors for Localized Clear Cell Renal Cell Carcinoma and Their Application in Adjuvant Therapy. Cancers 14, 239 (2022).

6. Flanigan, R. C., Campbell, S. C., Clark, J. I. & Picken, M. M. Metastatic renal cell carcinoma. Curr. Treat. Options Oncol. 4, 385–390 (2003).

7. Duran, I. et al. Resistance to Targeted Therapies in Renal Cancer: The Importance of Changing the Mechanism of Action. Target. Oncol. 12, 19–35 (2017).

8. Drugs Approved for Kidney Cancer - NCI. https://www.cancer.gov/about-cancer/treatment/drugs/kidney (2011).

9. Motzer, R. J. et al. Nivolumab versus Everolimus in Advanced Renal-Cell Carcinoma. N. Engl. J. Med. 373, 1803–1813 (2015).

10. Hartmann, J. T. & Bokemeyer, C. Chemotherapy for renal cell carcinoma. Anticancer Res. 19, 1541–1543 (1999).

11. Santos, M. L. C., de Brito, B. B., da Silva, F. A. F., Botelho, A. C. D. S. & de Melo, F. F. Nephrotoxicity in cancer treatment: An overview. World J. Clin. Oncol. 11, 190–204 (2020).

12. Wang, Y., Chen, X., Tang, N., Guo, M. & Ai, D. Boosting Clear Cell Renal Carcinoma-Specific Drug Discovery Using a Deep Learning Algorithm and Single-Cell Analysis. Int. J. Mol. Sci. 25, 4134 (2024).

13. Li, X. et al. Prediction of drug candidates for clear cell renal cell carcinoma using a systems biology-based drug repositioning approach. EBioMedicine 78, 103963 (2022).

14. Turanli, B. et al. Systems biology based drug repositioning for development of cancer therapy. Semin. Cancer Biol. 68, 47–58 (2021).

15. Li, R. et al. Mapping single-cell transcriptomes in the intra-tumoral and associated territories of kidney cancer. Cancer Cell 40, 1583–1599.e10 (2022).

16. Zhang, Y. et al. Single-cell analyses of renal cell cancers reveal insights into tumor microenvironment, cell of origin, and therapy response. Proc. Natl. Acad. Sci. U. S. A. 118, e2103240118 (2021).

17. Zhang, M. et al. Single cell analysis reveals intra-tumour heterogeneity, microenvironment and potential diagnosis markers for clear cell renal cell carcinoma. Clin. Transl. Med. 12, e713 (2022).

18. Zvirblyte, J. et al. Single-cell transcriptional profiling of clear cell renal cell carcinoma reveals a tumor-associated endothelial tip cell phenotype. *Commun*. Biol. 7, 1–15 (2024).

19. Mei, S. et al. Single-cell analysis of immune and stroma cell remodeling in clear cell renal cell carcinoma primary tumors and bone metastatic lesions. Genome Med. 16, 1 (2024).

20. Creighton, C. J. et al. Comprehensive molecular characterization of clear cell renal cell carcinoma. Nature 499, 43–49 (2013).

21. Laskar, R. S. et al. Sexual dimorphism in cancer: insights from transcriptional signatures in kidney tissue and renal cell carcinoma. Hum. Mol. Genet. 30, 343–355 (2021).

22. Chawla, N. V., Bowyer, K. W., Hall, L. O. & Kegelmeyer, W. P. SMOTE: Synthetic Minority Over-sampling Technique. J. Artif. Intell. Res. 16, 321–357 (2002).

23. Ke, G., et al. LightGBM: A Highly Efficient Gradient Boosting Decision Tree.

24. Hicks, S. A. et al. On evaluation metrics for medical applications of artificial intelligence. Sci. Rep. 12, 5979 (2022).

25. Szklarczyk, D. et al. The STRING database in 2023: protein-protein association networks and functional enrichment analyses for any sequenced genome of interest. Nucleic Acids Res. 51, D638–D646 (2023).

26. Han, N. et al. Identification of SARS-CoV-2-induced pathways reveals drug repurposing strategies. Sci. Adv. 7, eabh3032 (2021).

27. Guney, E., Menche, J., Vidal, M. & Barábasi, A.-L. Network-based in silico drug efficacy screening. Nat. Commun. 7, 10331 (2016).

28. Tsherniak, A. et al. Defining a Cancer Dependency Map. Cell 170, 564–576.e16 (2017).

29. Thul, P. J. et al. A subcellular map of the human proteome. Science 356, eaal3321 (2017).

30. Heyn, P., Kalinka, A. T., Tomancak, P. & Neugebauer, K. M. Introns and gene expression: cellular constraints, transcriptional regulation, and evolutionary consequences. BioEssays News Rev. Mol. Cell. Dev. Biol. 37, 148–154 (2015).

31. STRING: functional protein association networks. https://string-db.org/.

32. Shuch, B., Linehan, W. M. & Srinivasan, R. Aerobic glycolysis: a novel target in kidney cancer. Expert Rev. Anticancer Ther. 13, 711–719 (2013).

33. Yong, C., Stewart, G. D. & Frezza, C. Oncometabolites in renal cancer. Nat. Rev. Nephrol. 16, 156–172 (2020).

34. Linehan, W. M., Srinivasan, R. & Schmidt, L. S. The genetic basis of kidney cancer: a metabolic disease. Nat. Rev. Urol. 7, 277–285 (2010).

35. Chakraborty, S., Balan, M., Sabarwal, A., Choueiri, T. K. & Pal, S. Metabolic reprogramming in renal cancer: Events of a metabolic disease. Biochim. Biophys. Acta Rev. Cancer 1876, 188559 (2021).

36. Zerbini, L. F. et al. Computational repositioning and preclinical validation of pentamidine for renal cell cancer. Mol. Cancer Ther. 13, 1929–1941 (2014).

37. Rausch, M. et al. Drug Repurposing to Identify a Synergistic High-Order Drug Combination to Treat Sunitinib-Resistant Renal Cell Carcinoma. Cancers 13, 3978 (2021).

38. Feng, X., Zhang, L., Tu, W. & Cang, S. Frequency, incidence and survival outcomes of clear cell renal cell carcinoma in the United States from 1973 to 2014: A SEER-based analysis. Medicine (Baltimore) 98, e16684 (2019).

39. Yu, C.-P. et al. Estrogen inhibits renal cell carcinoma cell progression through estrogen receptor-β activation. PloS One 8, e56667 (2013).

40. Pizzocaro, G. et al. Adjuvant Medroxyprogesterone Acetate to Radical Nephrectomy in Renal Cancer: 5-Year Results of a Prospective Randomized Study. J. Urol. (1987) doi:10.1016/S0022-5347(17)43647-0.

41. Gao, Y., Ding, Y., Tai, X., Zhang, C. & Wang, D. Ponatinib: An update on its drug targets, therapeutic potential and safety. Biochim. Biophys. Acta BBA - Rev. Cancer 1878, 188949 (2023).

42. Sager, R. A. et al. Therapeutic potential of CDK4/6 inhibitors in renal cell carcinoma. Nat. Rev. Urol. 19, 305–320 (2022).

43. Elizarraras, J. M. et al. WebGestalt 2024: faster gene set analysis and new support for metabolomics and multi-omics. Nucleic Acids Res. 52, W415–W421 (2024).

44. Hainsworth, J. D. et al. Treatment of metastatic renal cell carcinoma with a combination of bevacizumab and erlotinib. J. Clin. Oncol. Off. J. Am. Soc. Clin. Oncol. 23, 7889–7896 (2005).

45. Bridgeman, V. L. et al. Preclinical Evidence That Trametinib Enhances the Response to Antiangiogenic Tyrosine Kinase Inhibitors in Renal Cell Carcinoma. Mol. Cancer Ther. 15, 172–183 (2016).

46. Czarnecka, A. M., Niedzwiedzka, M., Porta, C. & Szczylik, C. Hormone signaling pathways as treatment targets in renal cell cancer (Review). Int. J. Oncol. 48, 2221–2235 (2016).

47. Huynh, T. P., et al. Glucocorticoids suppress renal cell carcinoma progression by enhancing Na, K-ATPase beta-1 subunit expression. PloS One 10, e0122442 (2015).

