## Supplementary Files for "A systems-level machine learning approach uncovers therapeutic targets in clear cell renal cell carcinoma"

### Supplementary Files Information

- Supplementary File 1: List of all genes with AUC
- Supplementary File 2: Log2 fold change in expression in tumour cells compared to non-tumour cells
- Supplementary File 3: List of all key proteins as identified by the network analysis
- Supplementary File 4: Results of drug proximity for FDA approved drugs
- Supplementary File 5: Results of drug proximity for non-FDA approved drugs

| Dataset | Accuracy | Precision | Recall | F1 Score | AUC |
| --- | --- | --- | --- | --- | --- |
| American | 0.97 | 0.99 | 0.96 | 0.97 | 0.99 |
| Chinese | 0.97 | 0.98 | 0.98 | 0.98 | 0.99 |
| Lithuanian | 0.98 | 0.98 | 1.00 | 0.99 | 0.99 |
| Metastasis | 0.99 | 0.99 | 1.00 | 1.00 | 0.92 |

**Supplementary Table 1: Performance of tumour classification using 96 drug target genes across external datasets** Summary of classification performance (accuracy, precision, recall, F1 score, and AUC) for the binary LightGBM model using the initial 96-gene target set. The classifier was evaluated on four independent scRNA-seq datasets from the United States, China, Lithuania, and a ccRCC metastatic cohort.

| Gene name | Expression | Function in cancer |
| --- | --- | --- |
| ENO1 | upregulated | Glycolytic enzyme, has been implicated in metabolic reprogramming, immune evasion and apoptosis inhibition^1^ |
| CD52 | downregulated | CD52 is a glycoprotein found on the surface of immune cells that has been shown to regulate immune invasion^2^ |
| PTPRC | downregulated | Inhibits JAK family kinases^3^ |
| SPP1 | upregulated | Induction of epithelial-mesenchymal transition and drug resistance^4^ |
| DAB2 | upregulated | Loss is associated with MAPK, Wnt and TGFβ signalling that drives tumour progression^5^ |
| HINT1 | upregulated | Potentially limits CD4+ T cell infiltration^6^ |
| HLA-E | downregulated | Negative immunomodulation by overexpression^7^ |
| TIMP1 | upregulated | Promotes tumorigenesis via epithelial-mesenchymal transition in ccRCC^8^ |
| PGK1 | upregulated | Promotes tumorigenesis and sorafenib resistance in ccRCC^9^ |
| CLU | upregulated | Potential driver of metastasis in ccRCC^10^ |
| CFL1 | downregulated | Regulator of B and T cell response^11^ |
| CCND1 | upregulated | Promotes cell division and has been implicated in RCC progression^12^ |
| CRYAB | upregulated | Upregulation leads to poor outcomes in ccRCC^13^ |
| VIM | upregulated | Induction of epithelial-mesenchymal transition^14^ |
| KRT8 | upregulated | Promotes metastasis and STAT3 signalling^15^ |
| KRT18 | upregulated | Promotes cancer progression through MAPK signalling^16^ |
| CD63 | upregulated | Promotes metastasis and alters platelet signalling^17^ |
| PEBP1 | upregulated | Low expression promotes metastasis in ccRCC^18^ |
| SERPINA1 | upregulated | Promotes proliferation and migration of ccRCC^19^ |
| TPM1 | upregulated | Tropomyosin-1 promotes cancer cell apoptosis via the p53-mediated mitochondrial pathway in renal cell carcinoma^20^ |
| CST3 | downregulated | Immune regulator^21^ |
| FTL | upregulated | Potentially promotes iron retention which supports tumour growth^22^ |
| RAC2 | downregulated | Promotes ccRCC progression^23^ |

**Supplementary Table 2: Functional characterization of refined drug targets in ccRCC**
Detailed annotation of selected drug target genes following network refinement. Includes expression status (up/downregulated), functional relevance in cancer, and literature support. Bolded genes indicate inclusion in the final drug proximity analysis. Genes are sorted by descending single-feature AUC values.

| Dataset | Accuracy | Precision | Recall | F1 Score | AUC |
| --- | --- | --- | --- | --- | --- |
| American | 0.76 | 0.97 | 0.57 | 0.72 | 0.95 |
| Chinese | 0.91 | 0.98 | 0.92 | 0.95 | 0.96 |
| Lithuanian | 0.98 | 0.98 | 1.00 | 0.99 | 0.99 |
| Metastasis | 0.91 | 0.99 | 0.92 | 0.95 | 0.62 |

**Supplementary Table 3: Performance of tumour classification using the refined 17-gene target set across external datasets** Summary of classification performance (accuracy, precision, recall, F1 score, and AUC) for the LightGBM model using the proximity-refined 17-gene target signature. The classifier was evaluated on four independent scRNA-seq datasets from the United States, China, Lithuania, and a ccRCC metastatic cohort.

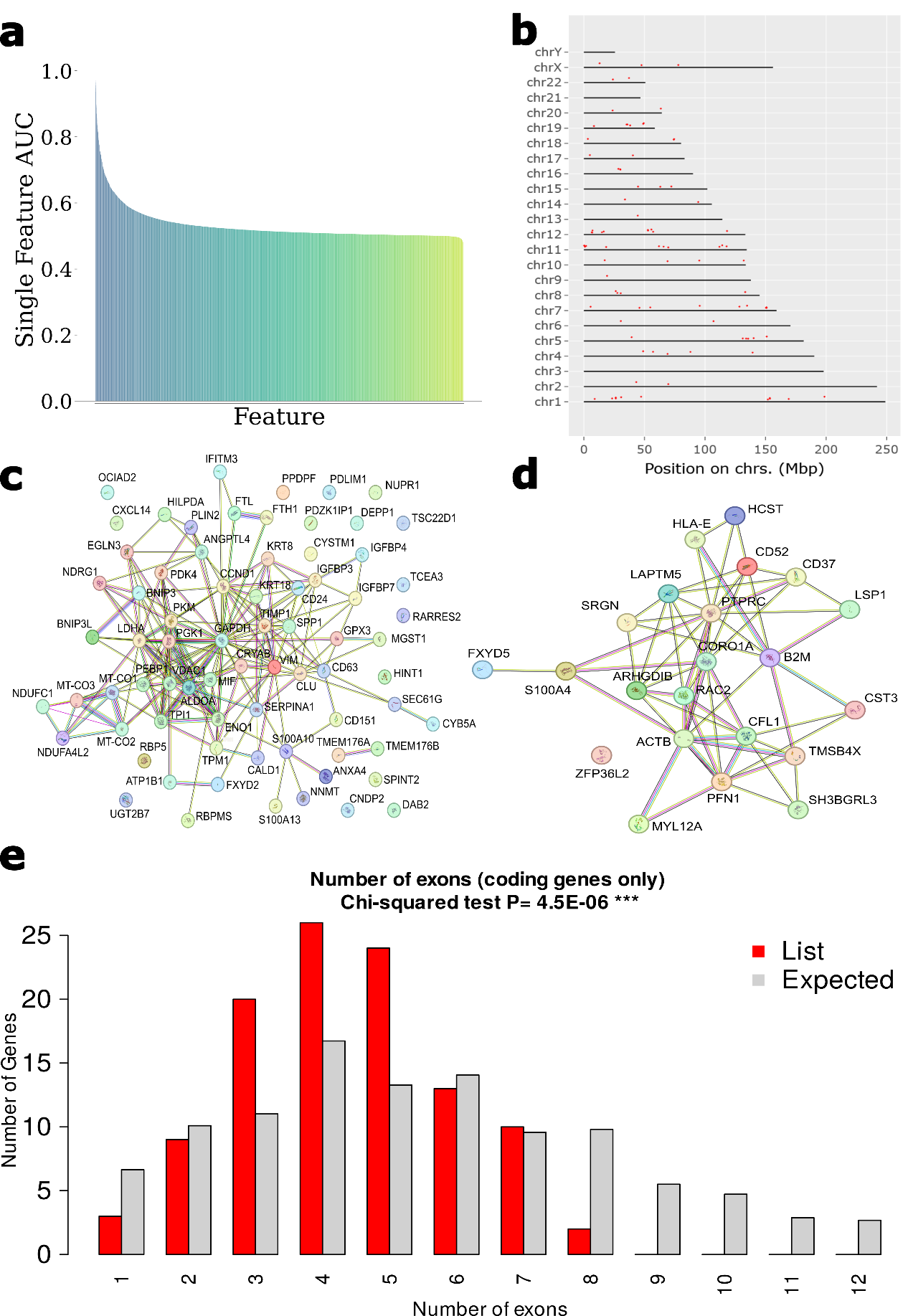

**Supplementary Figure 1: Extended characterization of drug target features.**
a) Single-feature AUC analysis reveals a subset of genes with strong individual discriminatory power (AUC > 0.8), supporting their use in tumour–normal classification. b) Chromosomal distribution indicates wide genomic spread, with no dominant chromosomal clustering. c–d) Protein–protein interaction (PPI) networks of upregulated (c) and downregulated (d) genes show higher connectivity among upregulated targets, suggesting coordinated activation of pro-tumorigenic pathways. e) Target genes exhibit a shorter exon count on average compared to the genome-wide background, consistent with the compact architecture of genes involved in regulatory and signal transduction functions.

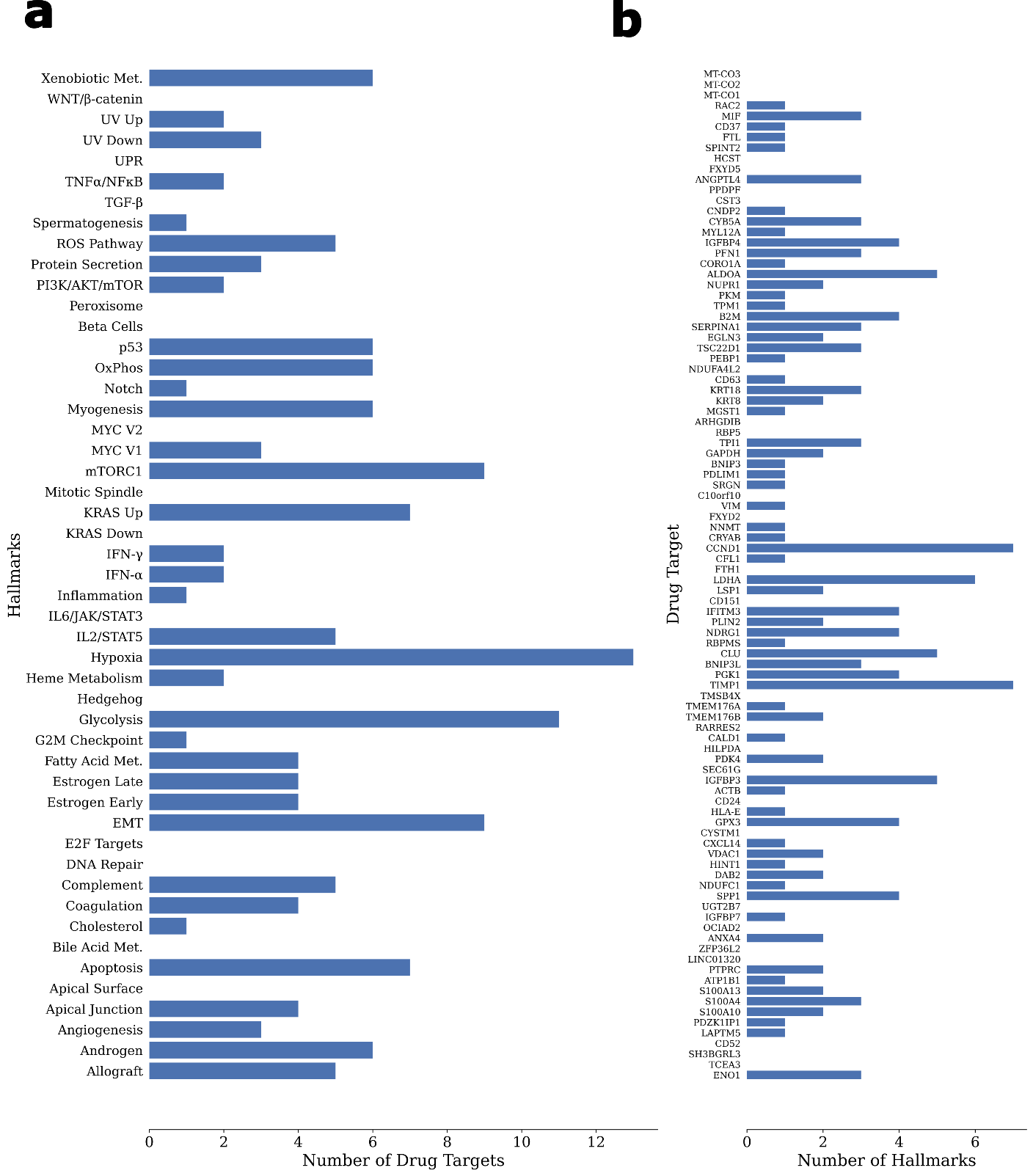

**Supplementary Figure 2: Association of drug targets with cancer hallmark pathways.**
Gene set enrichment analysis shows that selected targets are associated with multiple MSigDB hallmark categories, including proliferation, immune evasion, and metabolism. a) Target genes are most enriched in hallmarks related to epithelial–mesenchymal transition, glycolysis, and angiogenesis, aligning with known ccRCC biology. b) The majority of selected targets are linked to multiple hallmarks, highlighting their roles in tumour progression.

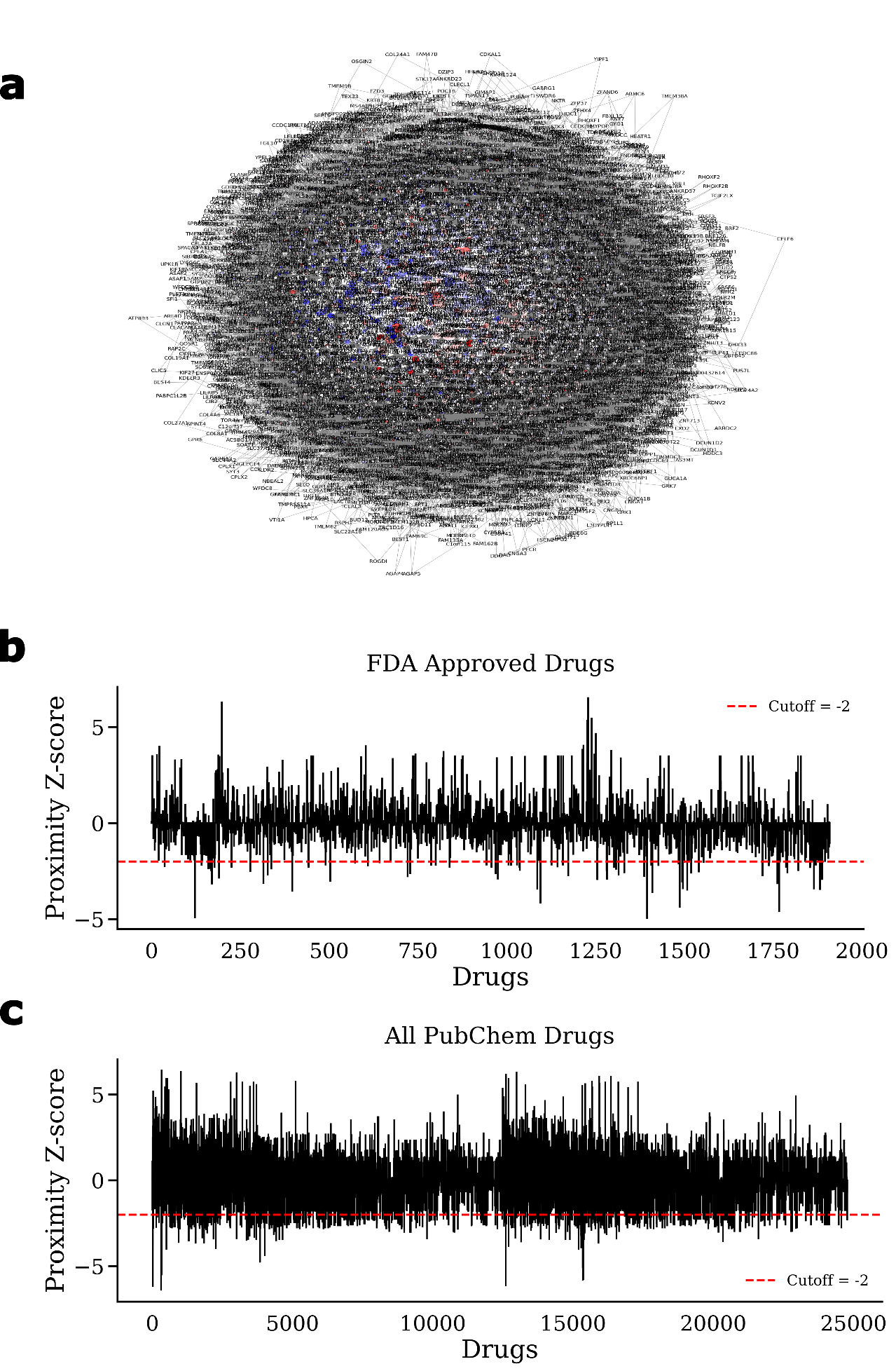

**Supplementary Figure 3: Extended** **Network analysis for drug target refinement and hit identification** a) Full gene interaction network, with upregulated genes shown in red, downregulated genes in blue, and genes with no notable changes in white. b) Z-proximity scores of FDA approved drugs to the refined drug targets with the indicated cutoff of -2. d) Drug proximity analysis of PubChem compounds with identical proximity cutoff.

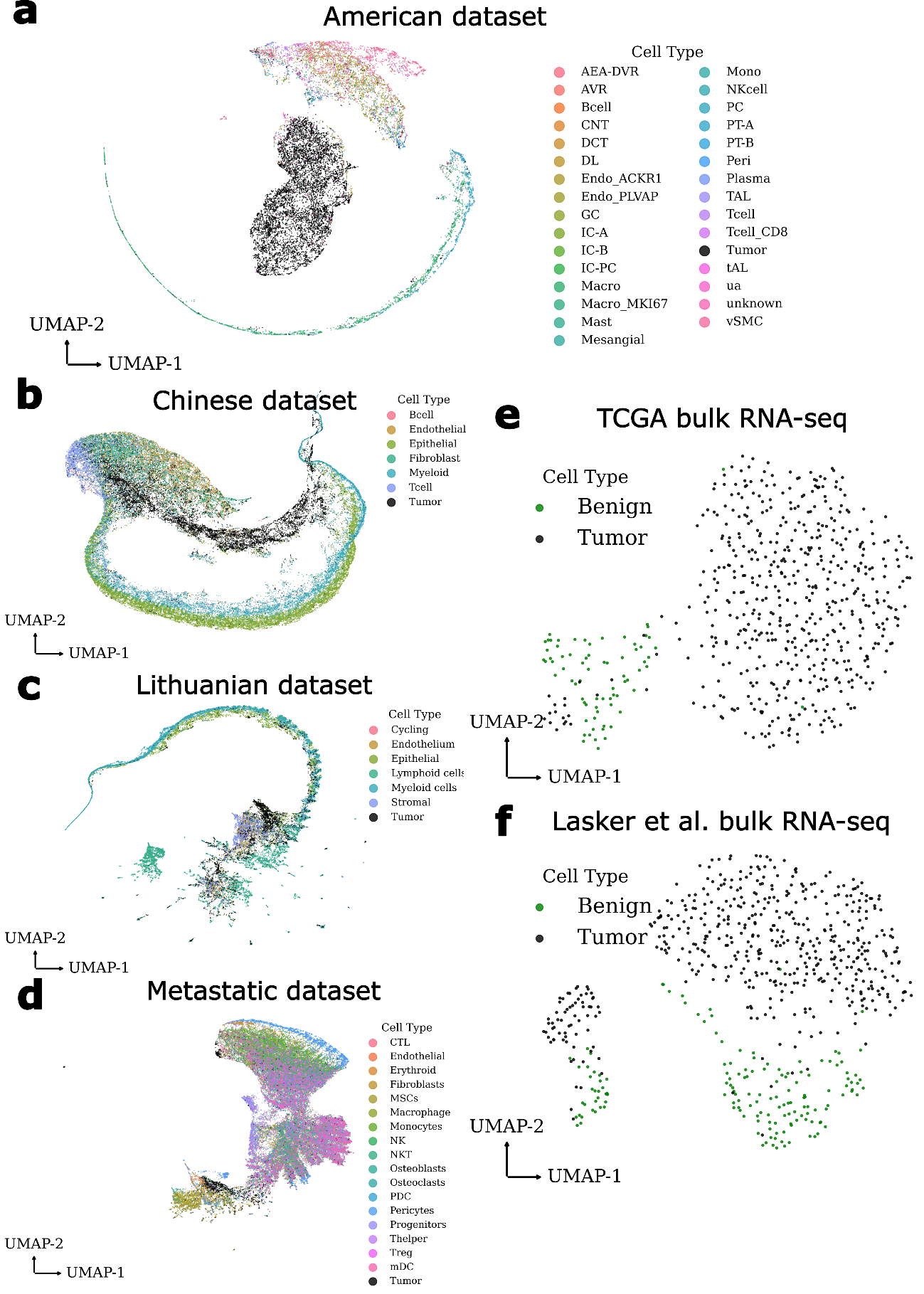

**Supplementary Figure 4: UMAP-based validation of refined drug targets across external datasets.**
UMAP projections demonstrate that the refined 17-genes robustly separate tumour and non-tumour cells across diverse datasets. a–d) Clear separation is observed in single-cell datasets from: the United States (Zhang et al. 2021) (**a**), China (Zhang et al. 2022) (**b**), Lithuania (Zvirblyte et al. 2024) (**c**), and ccRCC bone metastases (Mei et al. 2024) d). e–f) Even in bulk RNA-seq datasets, where cellular signals are averaged, the signature retains discriminative power: TCGA KIRC (e) and an independent U.S. bulk cohort (Laskar et al.) f), highlighting the translational robustness of the selected gene set.

### References

1.	Huang, C. K., Sun, Y., Lv, L. & Ping, Y. ENO1 and Cancer. *Mol. Ther. Oncolytics* **24**, 288–298 (2022).

2.	Wang, J. *et al.* CD52 Is a Prognostic Biomarker and Associated With Tumor Microenvironment in Breast Cancer. *Front. Genet.* **11**, 578002 (2020).

3.	Porcu, M. *et al.* Mutation of the receptor tyrosine phosphatase PTPRC (CD45) in T-cell acute lymphoblastic leukemia. *Blood* **119**, 4476–4479 (2012).

4.	Matsubara, E. *et al.* The Significance of SPP1 in Lung Cancers and Its Impact as a Marker for Protumor Tumor-Associated Macrophages. *Cancers* **15**, 2250 (2023).

5.	Price, Z. K. *et al.* Disabled-2 (DAB2): A Key Regulator of Anti- and Pro-Tumorigenic Pathways. *Int. J. Mol. Sci.* **24**, 696 (2022).

6.	Wang, X., Zhou, M. & Jiang, L. The oncogenic and immunological roles of histidine triad nucleotide-binding protein 1 in human cancers and their experimental validation in the MCF-7 cell line. *Ann. Transl. Med.* **11**, 147–147 (2023).

7.	Seliger, B. *et al.* HLA-E expression and its clinical relevance in human renal cell carcinoma. *Oncotarget* **7**, 67360–67372 (2016).

8.	Shou, Y. *et al.* TIMP1 Indicates Poor Prognosis of Renal Cell Carcinoma and Accelerates Tumorigenesis via EMT Signaling Pathway. *Front. Genet.* **13**, 648134 (2022).

9.	He, Y. *et al.* Novel inhibitors targeting the PGK1 metabolic enzyme in glycolysis exhibit effective antitumor activity against kidney renal clear cell carcinoma in vitro and in vivo. *Eur. J. Med. Chem.* **267**, 116209 (2024).

10.	Wang, X., Luo, L., Dong, D., Yu, Q. & Zhao, K. Clusterin plays an important role in clear renal cell cancer metastasis. *Urol. Int.* **92**, 95–103 (2014).

11.	Xing, J. *et al.* The role of actin cytoskeleton CFL1 and ADF/cofilin superfamily in inflammatory response. *Front. Mol. Biosci.* **11**, (2024).

12.	Karim, S. *et al.* Cyclin D1 as a therapeutic target of renal cell carcinoma- a combined transcriptomics, tissue microarray and molecular docking study from the Kingdom of Saudi Arabia. *BMC Cancer* **16**, 741 (2016).

13.	Ren, H. *et al.* CRYAB is upregulated and predicts clinical prognosis in kidney renal clear cell carcinoma. *IUBMB Life* **77**, e2938 (2025).

14.	Usman, S. *et al.* Vimentin Is at the Heart of Epithelial Mesenchymal Transition (EMT) Mediated Metastasis. *Cancers* **13**, 4985 (2021).

15.	Tan, H.-S. *et al.* KRT8 upregulation promotes tumor metastasis and is predictive of a poor prognosis in clear cell renal cell carcinoma. *Oncotarget* **8**, 76189–76203 (2017).

16.	Wang, P.-B., Chen, Y., Ding, G.-R., Du, H.-W. & Fan, H.-Y. Keratin 18 induces proliferation, migration, and invasion in gastric cancer via the MAPK signalling pathway. *Clin. Exp. Pharmacol. Physiol.* **48**, 147–156 (2021).

17.	Dudiki, T. *et al.* Mechanism of Tumor-Platelet Communications in Cancer. *Circ. Res.* **132**, 1447–1461 (2023).

18.	Yang, L. *et al.* Low expression of PEBP1P2 promotes metastasis of clear cell renal cell carcinoma by post-transcriptional regulation of PEBP1 and KLF13 mRNA. *Exp. Hematol. Oncol.* **11**, 87 (2022).

19.	Guo, L., An, T., Wan, Z., Huang, Z. & Chong, T. SERPINE1 and its co-expressed genes are associated with the progression of clear cell renal cell carcinoma. *BMC Urol.* **23**, 43 (2023).

20.	Tang, C. *et al.* Tropomyosin-1 promotes cancer cell apoptosis via the p53-mediated mitochondrial pathway in renal cell carcinoma. *Oncol. Lett.* **15**, 7060–7068 (2018).

21.	Zi, M. & Xu, Y. Involvement of cystatin C in immunity and apoptosis. *Immunol. Lett.* **196**, 80–90 (2018).

22.	Schnetz, M. *et al.* The Disturbed Iron Phenotype of Tumor Cells and Macrophages in Renal Cell Carcinoma Influences Tumor Growth. *Cancers* **12**, 530 (2020).

23.	Liu, Y. *et al.* RAC2 acts as a prognostic biomarker and promotes the progression of clear cell renal cell carcinoma. *Int. J. Oncol.* **55**, 645–656 (2019).

24.	GSEA | MSigDB | Human MSigDB Collections. https://www.gsea-msigdb.org/gsea/msigdb/human/collections.jsp#H.
